# ZIKA virus effects on neuroprogenitors are exacerbated by the main pyriproxyfen metabolite via thyroid hormone signaling disruption

**DOI:** 10.1101/2020.11.03.366468

**Authors:** Petra Spirhanzlova, Anthony Sébillot, Pieter Vancamp, Jean-David Gothié, Sébastien Le Mével, Michelle Leemans, Karn Wejaphikul, Marcel Meima, Bilal B. Mughal, Lucile Butruille, Pierre Roques, Sylvie Remaud, Jean-Baptiste Fini, Barbara A. Demeneix

## Abstract

North-Eastern Brazil saw intensive application of the insecticide pyriproxyfen (PPF) during the microcephaly outbreak caused by Zika virus (ZIKV). ZIKV requires the neural RNA-binding protein Musashi-1 to replicate. TH represses *MSI1*. Being a suspected TH disruptor, we hypothesized that co-exposure to the main metabolite of PPF, 4’-OH-PPF, would exacerbate ZIKV effects through increased MSI1 expression. This was tested using *in vitro* mouse neurospheres and an *in vivo* TH signaling reporter model, *Xenopus laevis*. TH signaling was decreased by 4’-OH-PPF in both models. In mouse-derived neurospheres the metabolite reduced neuroprogenitor proliferation as well as markers of neuronal differentiation. The results demonstrated that 4’-OH-PPF significantly induced MSI1 at both the mRNA and protein level, as well as *Fasn* mRNA. Other TH target genes were also significantly modified. Importantly, several key genes implicated in neuroprogenitor fate and commitment were not dysregulated by 4’-OH-PPF alone, but were in combination with ZIKV infection. These included the neuroprogenitor markers *Nestin, Egfr, Gfap, Dlx2* and *Dcx*. Unexpectedly, 4’-OH-PPF decreased ZIKV replication, although only at the fourth and last day of incubation, and RNA copy numbers stayed within the same order of magnitude. However, intracellular RNA content of neuroprogenitors was significantly decreased in the combined presence of the PPF metabolite and ZIKV. We conclude that 4’-OH-PPF interferes with TH action *in vivo* and *in vitro*, inhibiting neuroprogenitor proliferation. In the presence of ZIKV, TH signaling pathways crucial for cortical development are significantly impacted. This provides another example of viral effects that are exacerbated by drug or pesticide use.

**Significance statement:** In 2015, an increase in children born with unusually small heads (microcephaly) in North-Eastern Brazil was linked to infection with the ZIKA virus. An insecticide with thyroid hormone disruptive properties was used in the same areas. We investigated whether simultaneous exposure to the insecticide could increase viral susceptibility. The main metabolite 4’-OH-PPF dysregulated thyroid hormone signaling pathways crucial for brain development in both models used. Neural stem cells proliferated less and contained more Musashi-1, a protein the virus needs to replicate. Infecting stem cells pre-exposed to the endocrine disruptor did not amplify viral replication, but aggravated expression of genes implicated in brain development. Our results suggest the insecticide is particularly deleterious to brain development in areas with ZIKA virus prevalence.

## 1. Introduction

Multiple lines of evidence link infection by the ZIKA virus (ZIKV) to congenital Zika syndrome and the increased cases of neonatal microcephaly in Central and South America starting from early 2015 (1–5). Experiments on human brain organoids and neurospheres showed that neural stem cells (NSCs) are a direct target of ZIKV (6). ZIKV reduced proliferation, altered fate choice, increased cell death and decreased neuronal cell layer volumes (7, 8). ZIKV infection in mice led to smaller brains with thinner cortical layers and associated neurological disabilities by interfering with the cellular processes required for normal brain development (9).

Remarkably, a few regions in North-Eastern Brazil showed a much higher prevalence of ZIKV-associated microcephaly than others, prompting the question as to whether other factors could be linked to this regional effect (10, 11). It has been suggested that the use of the pesticide pyriproxyfen (PPF), a juvenile hormone analog, approved worldwide for use against household insects and in agriculture (12, 13), could be implicated in the rise of microcephaly cases (3, 14–16). High amounts of PPF were used in the same northeastern region where most ZIKV cases associated with microcephaly were found (17, 18). This could be yet another example of drug or pesticide exacerbating the effects of viral infection (19, 20). Even though a small cohort (< 80 cases) did not find a link between PPF and congenital microcephaly (3), it remains to be experimentally investigated whether or not the pesticide could be implicated in the microcephaly outbreak (15).

PPF was first introduced into Brazilian drinking water in late 2014 to control the *Aedes aegypti* mosquito population, the vector for the dengue and the ZIKA viruses. The WHO recommended that daily intake of PPF should not exceed 0.3 mg/L for an average adult, and advised that the concentration of PPF in drinking water containers should be lower than 0.01 mg/L (13). The main pathway of PPF metabolism is hydroxylation at the 4’-position, producing principally 4’-OH-PPF in both vertebrates and invertebrates (13, 21–23). PPF is documented to be rapidly taken up and metabolized, resulting in low bio-concentrations of the parent compound circulating in the organism (21). We therefore hypothesized that 4’-OH-PPF rather than PPF could be the main compound acting *in vivo*. We decided thus to carry out our experiments using the PPF main metabolite 4’-OH-PPF.

In 2017, Chavali et al. found that the RNA binding protein Musashi-1 (MSI1) is indispensable for ZIKV replication. MSI1 is an important translational regulator in NSCs in both vertebrates and invertebrates (24). Interestingly, we previously showed that this gene is downregulated *in vivo* in the murine subventricular zone (SVZ) by T_3_/thyroid hormone (TH) receptor α (TRα), when progenitors commit to the neuroblast lineage (25). THs regulate many brain developmental processes including cell proliferation, migration and differentiation to acquire the characteristic brain cyto-architecture (26, 27). In severe cases of hypothyroidism, disruption of these processes also leads to smaller brains and aberrant cortical layer formation (28). Considering that PPF was previously shown to interfere with thyroid-responsive endpoints in the amphibian metamorphosis assay and the male and female pubertal rat assays (29), we questioned if co-exposure to the two factors - 4’-OH-PPF and ZIKV - promoted increased viral replication.

We tested whether 4’-OH-PPF exposure could inhibit TH action, by increasing MSI1 expression and as such perturb NSC properties and replication. We used two TH model systems, *Xenopus laevis* tadpoles and adult mouse SVZ neuroprogenitors. In *Xenopus laevis*, we studied TH-disruptive effects *in vivo* using the XETA-assay, *msi1* expression as well as various morphometric and behavioral endpoints (30, 31). In mouse NSCs, we assessed *in vitro* effects on TH disruption, *Msi1* gene and protein expression as well as NSC proliferation. A final experimental approach examined ZIKV replication and TH target gene expression in mouse neuroprogenitors in the presence and absence of 4’-OH-PPF. Taken together, the results suggest that intense pesticide use could aggravate ZIKV infection by disrupting fetal brain developmen.

## 2. Results

### 4’-OH-PPF negatively affects TH signaling, tadpole motility and brain morphology

To investigate whether the PPF metabolite disrupted TH signaling, transgenic Tg(*thibz*:eGFP) tadpoles were exposed to an environmentally-relevant dose range of 4’-OH-PPF in the presence or absence of 5 nM T_3_ using the XETA assay. Control tadpole brains showed low basal fluorescence under physiological conditions (**Fig 1A, left panel**). As expected, adding 5 nM T_3_ strongly increased *eGFP* transcription, resulting in increased fluorescent activity (**Fig 1A, middle panel**). When simultaneously exposing to 4’-OH-PPF and T_3_, the fluorescent signal strongly decreased (**Fig 1A, right panel**). Testing the fluorescent signal in each condition with a specific concentration of 4’-OH-PPF + T_3_ showed the fluorescence in brains to be significantly reduced (**Fig 1B, C**, 0.05<p<0.0001). Further, even the lowest concentration (10^−7^ nM 4’-OH-PPF) reached significance (p<0.05) and the highest concentration, representing environmentally relevant levels had the strongest effects (**Fig 1C**, p<0.0001). These results suggest that 4’-OH-PPF exhibits TH-disrupting properties *in vivo*, and that short-term (72 h) exposure negatively impacts TR-mediated gene expression.

**Figure 1:**
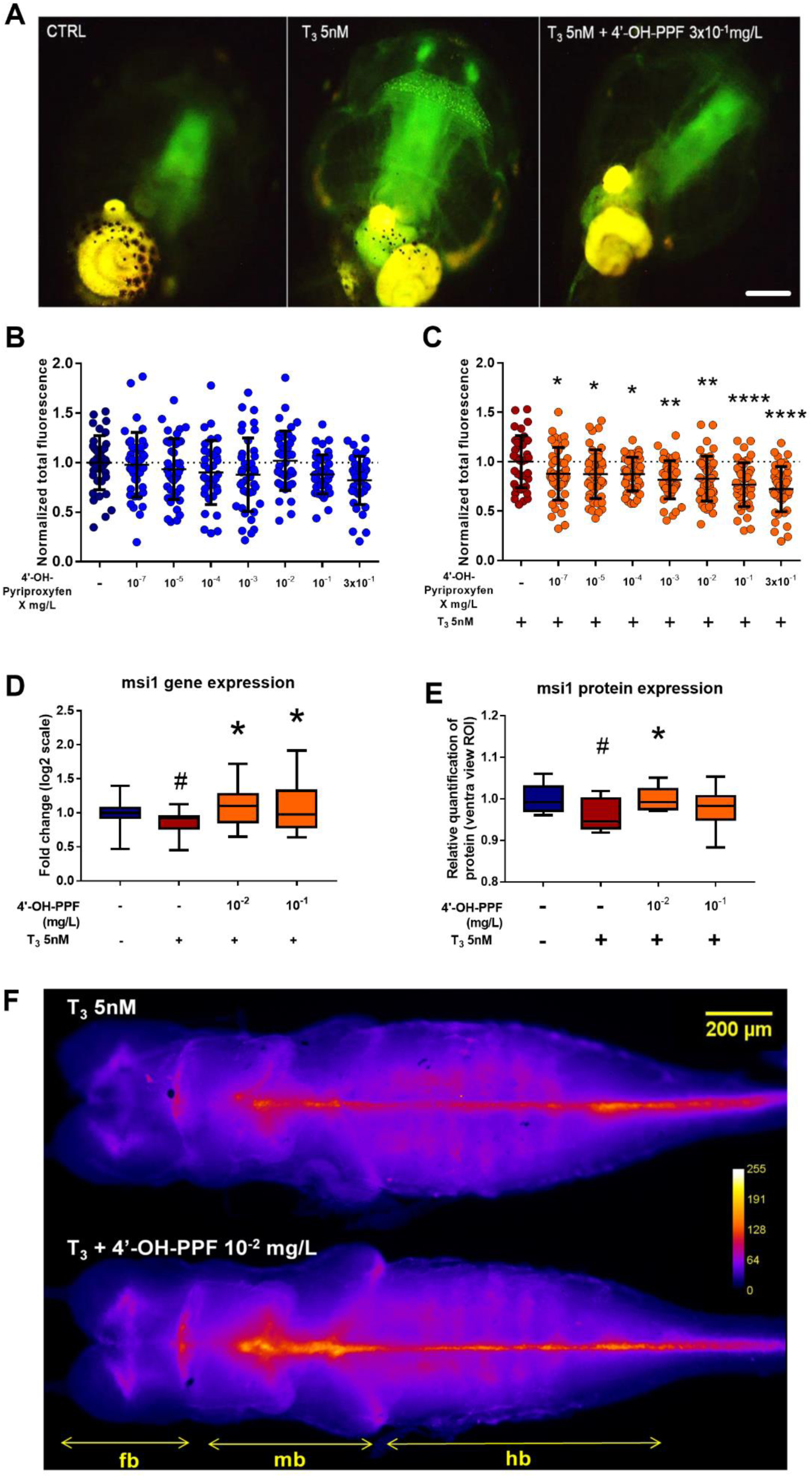
4’-OH-PPF affects thyroid hormone signaling, Musashi-1 gene expression and its encoded protein in *Xenopus laevis*. **(A)** Tg(*thibz*:eGFP) tadpoles exposed for 72 h to (from left to right) vehicle control, 5 nM T_3_ and 5 nM T_3_ + 3×10^−1^ mg/L 4’-OH-PPF. Scale bar: 500 μm. **(B, C)** 72 h exposure of NF45 Tg(*thibz*:eGFP) tadpoles to 4’-OH-PPF with **(B)** or without 5 nM T_3_ **(C)**. Values normalized to control group or T_3_-treated group, respectively (3 independent experiments; n=15 per experiment). **(D)** *msi1* gene expression in dissected brains from NF46 tadpoles exposed for 24 h to 4’-OH-PPF (10^−2^; 10^−1^ mg/L) together with 5 nM T_3_ (3 independent experiments, n=5 per experiment). **(E)** Msi1 protein expression in dissected brains from NF46 tadpoles exposed for 24 h to 4’-OH-PPF (10^−2^; 10^−1^ mg/L) together with 5 nM T_3_. Protein levels were quantified by measuring fluorescence intensity from dorsal views (2 independent experiments with 4-10 tadpoles per experiment). – Data were analyzed by a One-way ANOVA with Dunnett’s post-hoc test (Mean ± SEMs, * p<0.05, ** p<0.01, **** p <0.0001). Control and 5 nM T_3_ alone were compared with a - Mann-Whitney test (# p<0.05). **(F)** Representative pictures of dorsal views of NF46 *Xenopus* tadpoles exposed to 5 nM T_3_ or T_3_ + 10^−2^ mg/L 4’-OH-PPF and stained immunohistochemistically for Msi1. Msi1 levels are visualized as a heat map representing integrated density with arbitrary values from 0-256. Dark blue areas – no or low expression of Msi1; yellow/white – highest expression of Msi1); fb: forebrain, mb: midbrain, hb: hindbrain.

Experiments in JEG3 cells also showed TRα1-dependent luciferase expression to be affected at the highest dose used (3 mg/L 4’-OH-PPF) (**Fig S1A**, p<0.05). Similarly, applying the TR antagonist NH3 to the same cell line inhibited TRα1-dependent luciferase expression at the two highest concentrations tested (**Fig S1B**, p<0.05). This reemphasizes that high doses of the metabolite antagonize TR-dependent transcriptional activity.

We assessed whether 4’-OH-PPF affected tadpole behavior. Non-exposed tadpoles were active during the light periods and mostly inactive in the dark. Tadpoles exposed to 4’-OH-PPF exhibited reduced mobility compared to controls, reaching significance at the two highest concentrations (10^−1^ and 3×10^−1^ mg/L) (**Fig S2A, B**, p<0.05).

Next, total head size and brain areas of tadpoles were measured and brain/head ratios calculated. Tadpoles exposed to 3×10^−1^ mg/L 4’-OH-PPF had significantly smaller heads compared to controls (**Fig S2C**), corresponding to increased brain/head ratios (**Fig S2D**). However, the overall brain size of these tadpoles was not affected (**Fig S2E**). The lowest concentration of 10^−5^ mg/L 4’-OH-PPF significantly increased the width of forebrain and midbrain junction (**Fig S2F**, p<0.0001) and the width of the midbrain (**Fig S1G**, p<0.0001). These measurements collectively show that developmental exposure to 4’-OH-PPF leads to disproportionate brain dimensions.

### 4’-OH-PPF affects brain musashi-1 gene and protein expression in *Xenopus laevis* tadpoles

Next, we studied the consequences of a 24 h exposure to 4’-OH-PPF on musashi-1 (*msi1*) expression and its encoded protein in the brains of NF45 *Xenopus laevis* tadpoles *in vivo*. T_3_ significantly reduced *msi1* expression (**Fig 1D**, p<0.05) as well as the encoded protein (**Fig 1E**, p<0.05). At both concentrations used, 4’-OH-PPF increased *msi1* expression (p<0.05) in the presence of 5 nM T_3_ (**Fig 1D**, p<0.05), and significantly increased Msi1 protein levels at concentrations of 10^−2^ mg/L (**Fig 1E, F**, p<0.05). Moreover, Msi1 protein levels were predominantly upregulated in regions bordering the ventricular system, where most NSCs reside producing the cells of the central nervous system (**Fig 1F**). Hence, short exposure to 4’-OH-PPF at varying concentrations, and in presence of T_3_, increases Msi-1 levels *in vivo*, a crucial protein required for viral replication.

### 4’-OH-PPF ± T_3_ reduces proliferation of adult mouse NSCs

We next addressed whether 4’-OH-PPF ± 10 nM T_3_ altered the *in vitro* proliferation of primary neurospheres grown from adult mouse SVZ-NSCs, analyzing diameters and numbers of neurospheres (**Fig 2A**). After seven days, exposure to 10^−1^ and 3×10^−1^ mg/L 4’-OH-PPF reduced the average neurosphere diameter by 27.9% and 47.7% respectively, an effect also seen by adding T_3_ (**Fig 2B**, p<0.0001). Importantly, as neurospheres are 3D structures, the overall decrease in volume and cell number was greater. A diameter reduction of 28% corresponds to a volume decrease of 62.7% (ctrl: 641.4 mm^3^ vs 10^−1^ mg/L 4’-OH-PPF: 239.4 mm^3^), showing strongly impaired NSC proliferation following exposure to 4’-OH-PPF. Further, neurospheres exposed to the same concentrations of 4’-OH-PPF showed reduced growth capacity and a 70% reduction in neurosphere number (**Fig 2C**, p<0.0001), an effect seen with or without adding T_3_. All these results indicate that exposure to 4’-OH-PPF strongly impacts the capacity of neuroprogenitor proliferation.

**Figure 2.**
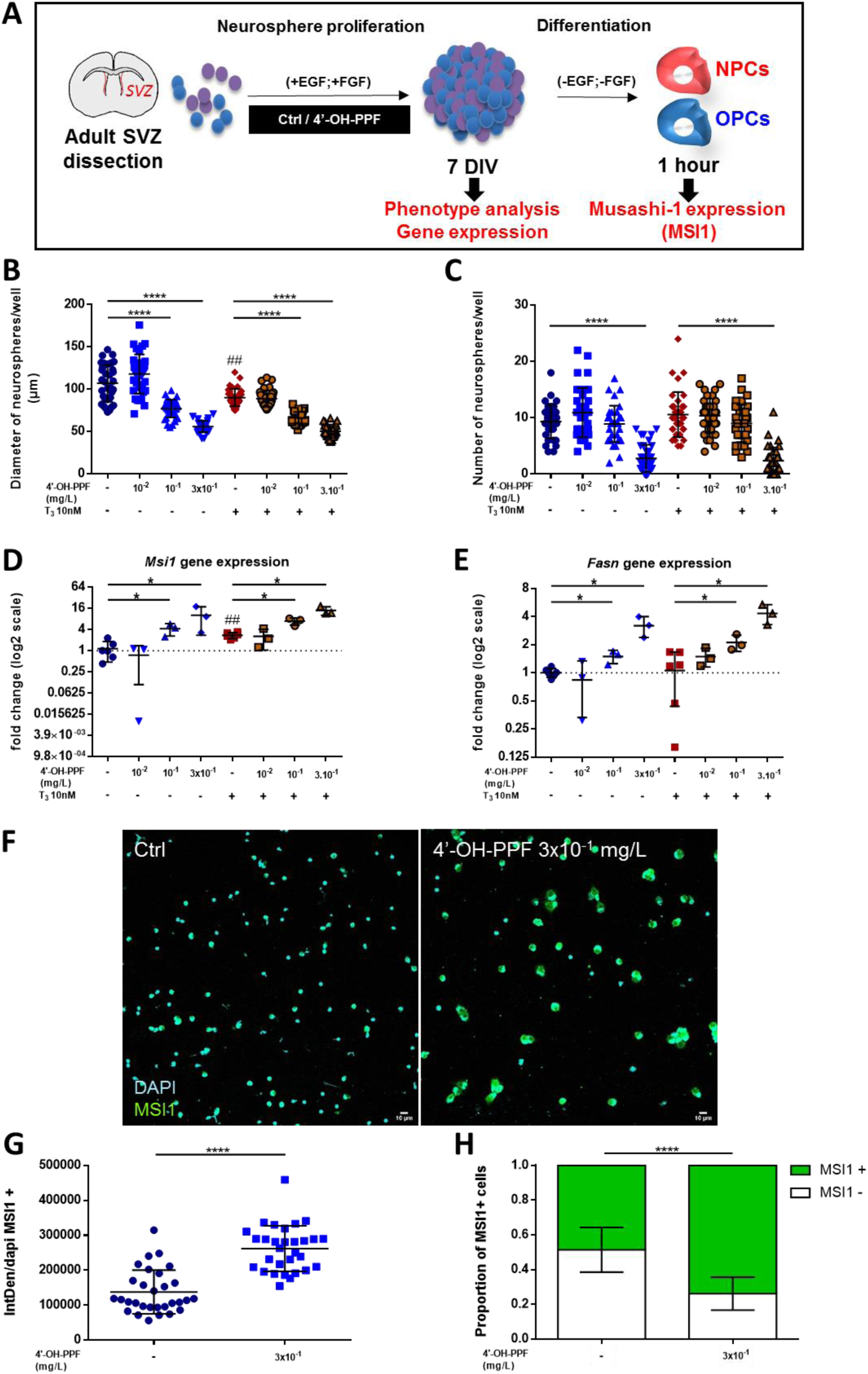
4’-OH-PPF exposure decreases NSCs proliferation and enhances *Fasn* expression levels, as well as *Msi1* mRNA and its encoded protein. **(A)** Schematic timeline illustrating the experimental design for the neurosphere assay. **(B)** Graphs showing neurosphere diameter in presence of 10 nM T_3_ ± 3×10^−1^ mg/L 4’-OH-PPF 10^−1^ versus controls without added T_3_ (n=12 wells/group, Kruskal-Wallis test: p<0.0001, followed by Mann-Whitney post-tests: ^##^: p=0.0011; ****: p<0.0001). **(C)** Graphs showing the average neurosphere number in presence of 10 nM T_3_ ± different doses of 4’-OH-PPF 10^−1^ mg/L versus controls without added T_3_ (n=12 wells/group, Kruskal-Wallis test: p<0.0001, followed by Mann-Whitney post-tests: ****: p<0.0001). **(D-E)** Graphs showing *Msi1* and *Fasn* mRNA expression levels in presence of 10 nM T_3_ ± different doses of 4’-OH-PPF (n=3-6 wells per group, Kruskal-Wallis test: *Fasn*, p=0.0019, *Msi1*, p=0.0009, followed by Mann-Whitney post-tests: *Fasn* *: p=0.0238, *Msi1*\*: p=0.0238). **(F)** Representative images of immunostained MSI1 (green) protein in cultured DAPI-positive adult mouse NSCs (blue) following exposure to 0.01% DMSO (Ctrl) or 3×10^−1^ mg/L 4’-OH-PPF 10^−1^ mg/L for 7 days and subsequent differentiation for 1 h, MSI1 protein levels strongly increased after exposure to 3×10^−1^ mg/L 4’-OH-PPF **(right panel)**. Scale bars: 10 µm. **(G)** MSI1 levels were analyzed by quantifying the integrative density of the green signal of MSI1+ cells, **(H)** and by measuring the relative proportion of MSI1+ cells in the total DAPI+ cell population (n=3 replicates/groups, n=10 images/replicates, Mann-Whitney test ****: p<0.0001). All graphs depict means ± SDs. Ctrl: control, DIV: day *in vitro*, EGF: epidermal growth factor, FGF: fibroblast growth factor, NPCs: neural progenitor cells, NSCs: neural stem cells, OPCs: oligodendrocyte progenitor cells, SVZ: subventricular zone.

### Exposure to 4’-OH-PPF increases *Fasn, Msi* and MSI1 expression in NSCs

We analyzed whether a seven day 4’-OH-PPF exposure modified *Fasn* and *Msi1* gene expression as well as MSI1 protein levels, factors required for NSC proliferation, using the same protocol as in **Fig 2A**. In mouse neurosphere cultures, environmentally relevant concentrations of 4’-OH-PPF ± 10 nM T_3_ induced a dose-dependent increase in *Msi1* expression (**Fig 2D**). Without adding T_3_, the highest doses of 4’-OH-PPF (range: 10^−1^ – 3.10^−1^ mg/L) induced *Msi1* expression 4.5-fold to 9.4-fold compared to controls (p<0.05). With the addition of T_3_, the induction was respectively 2.7-fold to 4.4-fold (**Fig 2D**, p<0.05). The highest doses of 4’-OH-PPF also induced *Fasn* expression in a parallel manner to that of *Msi1*, whether T_3_ was added or not (**Fig 2E**, p<0.05). We next performed immunostaining in 1 h differentiated neurospheres to quantify MSI1 protein expression and the proportion of MSI1-positive cells. Seven-days exposure to 3×10^−1^ mg/L 4’-OH-PPF increased the signal (**Fig 2F**), doubling MSI1 protein levels against controls (**Fig 2G**, p<0.0001) and increased the proportion of MSI-1 positive cells (control: 48.6%; 4’-OH-PPF 10^−1^ mg/L: 73.9%) (**Fig 2H**, p<0.0001).

### ZIKV with 4’-OH-PPF decreases NSC RNA content and modifies neuroprogenitor markers

Based on the previous findings, we set up a protocol for testing whether ZIKV infection was modulated by 4’-OH-PPF. To establish the exposure conditions and the ZIKV infection period, we carried out a preliminary experiment on murine neurospheres (**Fig S3**) to determine the temporal *Msi1* expression patterns during and after 4’-OH-PPF exposure. We found that exposing neurospheres to 4’-OH-PPF 10^−1^mg/L for 2 days before and 4 days after dissociation slightly but significantly induced *Msi1* as well as the strongly T_3_-responsive gene *Krüppel-like factor 9* (*Klf9*) (**Fig S3**, *Msi1, Klf9*: p<0.01).

Based on this knowledge, we then tested ZIKV infection. NSCs proliferated for a week and were exposed to 4’-OH-PPF for the last 5 days. We then dissociated the neurospheres and infected them with ZIKV for 4 days ± 4’-OH-PPF (**Fig 3A**). We measured ZIKV replication in two independent experiments, where we quantified RNA copy number (copies/mL) in the supernatant by RT-PCR each day post-infection. Viral replication increased as a function of time post-infection ± 4’-OH-PPF, increasing from 10^4^ to 10^8^ copies/mL. The presence of 4’-OH-PPF only significantly reduced replication at day 4 after ZIKV infection, but the values remained within the same order of magnitude (10^8^) (**Fig 3B**, p<0.0001 only at day 4).

**Figure 3:**
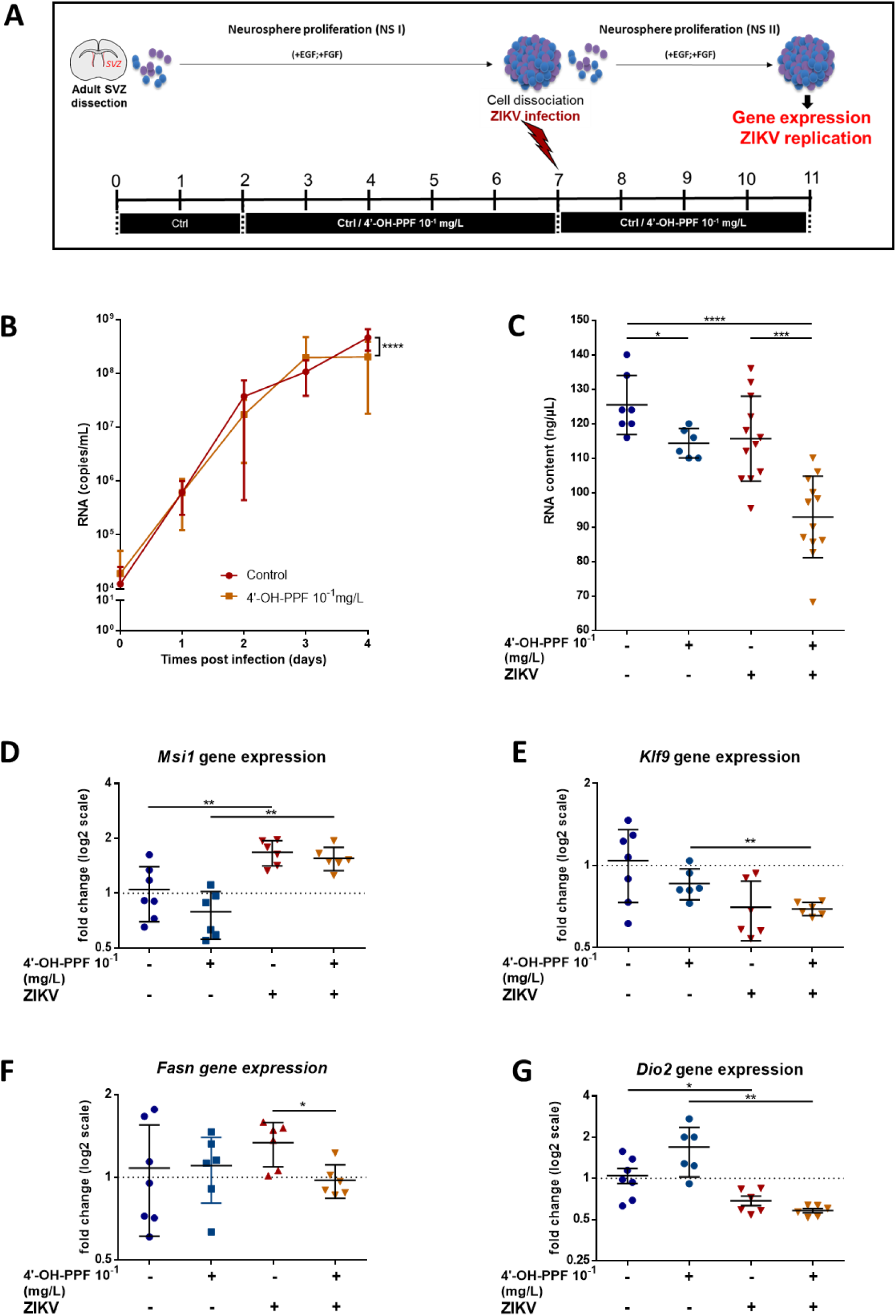
4’-OH-PPF exposure does not alter ZIKV replication in neurospheres, but decreases intracellular RNA content and dysregulates TH signaling. **(A)** Schematic timeline of the exposure and infection protocol. **(B)** ZIKV virus RNA copy number (copies/ml) at 0 and 4 days post-infection, showing the exponential increase over 4 days. Exposure to 4’-OH-PPF led to small but significant number of ZIKV particles at day 4, but no other day (2 independent experiments, n=6/group, 2-way ANOVA: p<0.0001, followed by Bonferroni’s multiple comparisons test: day 0-3: p>0.05, day 4: **** p<0.0001). **(C)** Graph showing neuroprogenitor intracellular RNA content (n= 6-12, Kruskal-Wallis test: p<0.0001, followed by Mann-Whitney post-tests: *: p=0.0105, ***: p=0.0002, ****: p<0.0001). **(D-G)** Graphs showing expression profiles of TH-target and neuroprogenitor marker genes in all four conditions (n=6-12). Kruskal-Wallis test: *Msi1*, p=0.012, *Klf9*, p=0.0306, *Fasn*, p=0.02055, *Dio2*, p=0.0009, followed by Mann-Whitney post-tests: *: p<0.05, **: p<0.01. For all graphs: n=7: Ctrl (no 4’-OH-PPF/no ZIKV), n=6: 4’-OH-PPF 10^−1^ mg/L (no ZIKV), n=12: ZIKV ± 4’-OH-PPF 10^−1^ mg/L. Graphs depict means ± SDs. Ctrl: control, EGF: epidermal growth factor, FGF: fibroblast growth factor, NSCs: neural stem cells, SVZ: subventricular zone, ZIKV: Zika virus.

We then analyzed the gene expression profile of well-known TH target genes at day 4 after ZIKV infection in the four conditions. These were with or without ZIKV and with or without 4’-OH-PPF. *Msi1* expression significantly increased in the presence of the ZIKV ± 10^−1^ mg/L 4’-OH-PPF (**Fig 3C**, control vs ZIKV, p<0.05; 4’-OH-PPF vs 4’-OH-PPF+ZIKV, p<0.001). *Msi1* expression was no longer induced after 9 days exposure to 4’-OH-PPF in absence of ZIKV, indicating continually fluctuating *Msi1* levels (**Fig. 3D**, left-hand scatter plots, p>0.05). However, ZIKV presence induced *Msi1* expression and the increase was maintained in the presence of 4’-OH-PPF (**Fig 3D**, right-hand scatter plots, p<0.01). The same observation applies to *Klf9* expression (**Fig 3E**). However, the presence of ZIKV in combination with 4’-OH-PPF significantly reduced *Klf9* levels (**Fig 3E**, 4’-OH-PPF vs 4’-OH-PPF + ZIKV, p<0.01) and *Fasn* expression (**Fig 3F**, p<0.05). The presence of 4’-OH-PPF induced a trend to increased *Dio2* expression (**Fig 3G**, left-hand scatter plots). Co-infecting with ZIKV reduced *Dio2* expression with or without 4’-OH-PPF (**Fig 3G**, right-hand scatter plots, p<0.05).

**Fig 4A** shows the markers expected at each stage of NSC progression. We analyzed *Gfap* as a general marker for NSCs, *Egfr, Nestin, Sox2* for activated NSCs, *Sox10* for oligodendocyte precursor cells, *Ng2* for immature oligodendrocytes, *Dlx2* for neuronal precursors, and *Dcx* for neuroblasts. *Gfap* levels were increased in the presence of ZIKV and even more when combined with 4’-OH-PPF (**Fig 4B**, p<0.01). *Egfr* and *Nestin* levels decreased as a function of viral presence, whereas with 4’-OH-PPF they increased significantly, but never returning to basal levels (**Fig 4C, D**, p<0.01 in all cases). *Sox2* and *Sox10* expression decreased with 4’-OH-PPF without ZIKV (**Fig 4E, F**, p<0.05 and p<0.01, respectively). *Ng2* expression was only slightly increased by ZIKV and 4’-OH-PPF exposure (**Fig 4G**, p<0.05). Both *Dlx2* and *Dcx* levels increased as a function of viral exposure (**Fig 4H, I**, p<0.01), but the presence of 4’-OH-PPF increased that of *Dlx2* (**Fig 4H**, p<0.05) whereas that of *Dcx* decreased (**Fig 4I**, p < 0.05).

**Figure 4:**
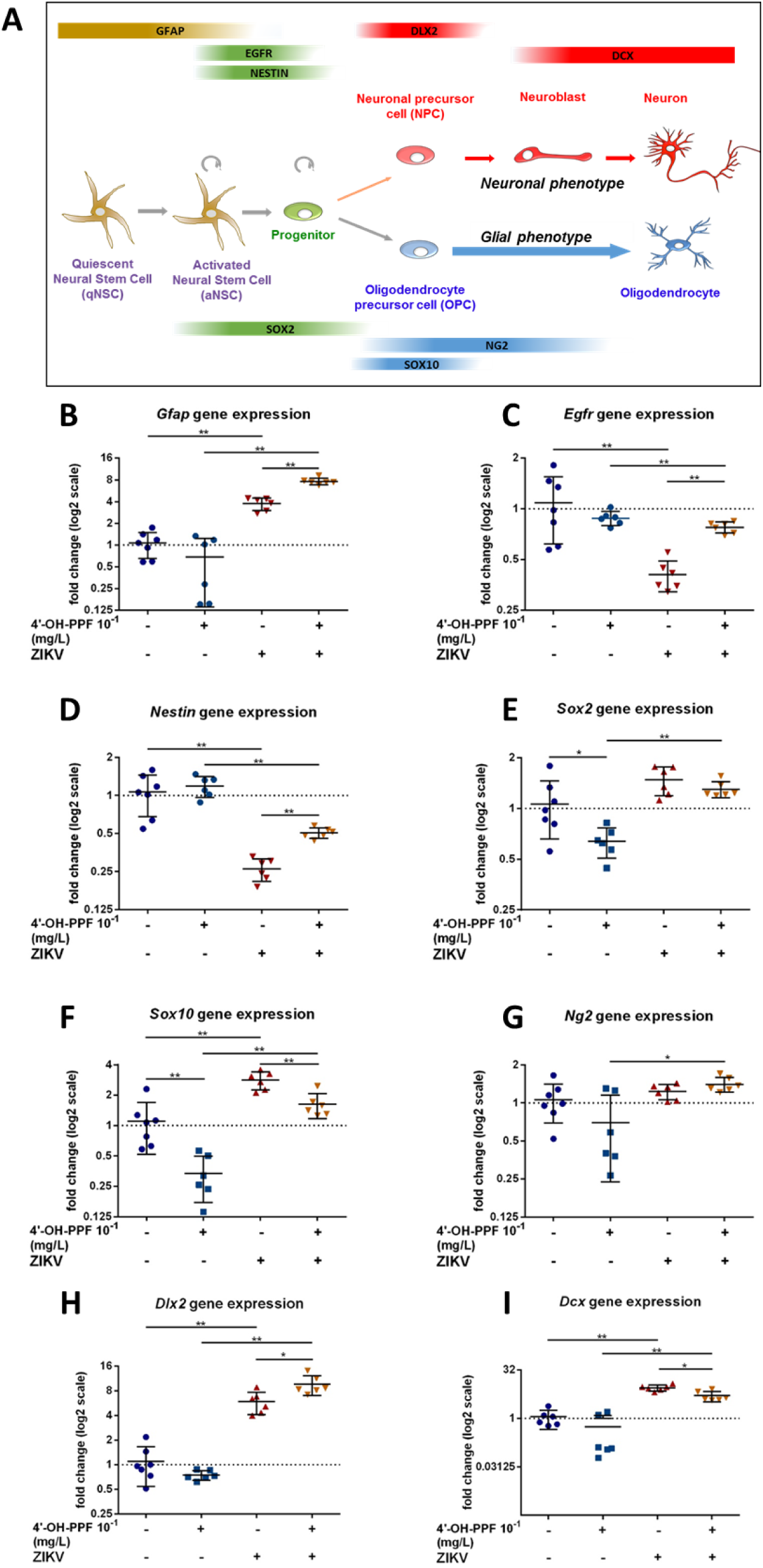
Key TH-target genes and those involved in NSC processes are modified by ZIKV infection with and without 4’-OH-PPF. **(A)** Scheme showing neuro- and gliogenesis in adult the subventricular zone and associated biomarkers during NSC commitment. **(B-I)** Graphs showing gene expression in NSCs: n=7: Ctrl (no 4’-OH-PPF/no ZIKV), n=6: 4’-OH-PPF 10^−1^ mg/L (no ZIKV), n=12: ZIKV ± 4’-OH-PPF 10^−1^ mg/L. Kruskal-Wallis test: *Gfap*, p=0.0001, *Egfr*, p=0.0017, *Sox2*, p=0.0028, *Ng2*, p=0.0212, *Nestin, Sox10, Dlx2, Dcx*, p=0.0002, followed by Mann-Whitney post-tests: *: p<0.05, **: p<0.01. Graphs depict means ± SDs. aNSC: activated neural stem cell, qNSC: quiescent neural stem cell, NPC: neural progenitor cells, NSCs: neural stem cells, OPC: oligodendrocyte progenitor cell.

## 3. Discussion

ZIKV infection was proposed as the causative factor in the sudden microcephaly outbreak in South and Central America in 2015. Data from human cultured NSCs, brain organoids and animal models indicate that these neuropathies predominantly resulted from a ZIKV-associated decrease in NSC proliferation, cell-cycle dysregulation, altered fate choice and increased apoptosis during brain development (7, 32–37). In the present study, we investigated whether there could be a connection between the pesticide PPF and the microcephaly outbreak in certain regions in North-Eastern Brazil where prevalence and use was much higher as compared to other regions (14, 15). PPF could increase virus susceptibility, aggravating the adverse effects ZIKV has on brain development.

Our hypothesis was based on two ideas. First, THs are key to evolutionary expansion of brain size and complexity, a primary characteristic of humans (38). These hormones regulate NSC pool expansion, progenitor multiplication, fate choice, migration and differentiation, important processes underlying normal brain development (27). Amongst the genes specifically implicated in human brain evolution are TH-regulated genes such as transthyretin (*TTR*) (38) and sonic hedgehog (*SHH*) (39). Many genes involved in neuroprogenitor proliferation have also undergone human-specific acceleration (39). One such gene is adenlyate-cyclase-activating-polypetide 1 (*ADCYAP1*), which is also a TH-regulated gene (40). All primary microcephaly genes are implicated in cell cycle or neuroprogenitor control (38). Second, classic cases of viral infection exacerbated by pesticide or drug use exist. Reye’s Syndrome has been linked to aspirin usage in children and viral meningitis (20) and vitamin deficiency to worsening of COVID-19 symptoms (19).

A central idea to our hypothesis was that the main metabolite of PPF, 4’-OH-PPF, might affect TH signaling and thereby increase expression of the negatively TH-regulated gene *Msi1*. As its encoded protein MSI1 is a crucial mediator of viral replication, the metabolite could exacerbate ZIKV replication. We tested our hypothesis in two models: *Xenopus laevis* tadpoles and *in vitro* mouse neurospheres. Lastly, we infected NSCs pre-exposed to 4’-OH-PPF with ZIKV to evaluate viral replication and effects on the TH signaling pathway.

### 4’-OH-PPF disrupts TH signaling in *Xenopus* tadpoles and mouse NSCs

The results from the XETA assay confirmed our initial hypothesis that the lowest environmentally relevant concentrations of 4’-OH-PPF exhibited anti-thyroid activity in the tadpole brain. Furthermore, we found that major brain compartments developed disproportionately, suggesting brain development was perturbed. This could also explain the observed decreased tadpole mobility.

We then extended these findings with an *in vitro* gene expression study on mouse NSCs. The metabolite modified TH-responsive gene expression, increasing *Msi1* and *Fasn*. The RNA binding protein *Msi1* is required for repression of p21^WAF-1^ protein implicated in NSC proliferation (41). Lipid synthesis requires *Fasn* (42). All membranes, including cell and mitochondrial structures, are dependent on correct amounts of *Fasn*. Exposing Jurkat, HeLa and trophoblast cells to PPF also disrupted membrane stability, lowering cell viability (43). Interestingly, PPF enhanced viral replication when simultaneously exposed to vesicular stomatitis virus (43).

We found that NSC proliferation was strongly decreased following 4’-OH-PPF exposure. Other genes include a tendency to increased *Dio2* expression and significantly decreased expression of the stem cell marker *Sox2*, as well as *Sox10*, both T_3_-negatively regulated genes. The latter directs NSC fate towards the oligodendroglial lineage (44). These observations could indicate decreased T_3_ levels in NSCs suggesting that they are in a local hypothyroid state. The results on JEG3 cells corroborated this concept, namely that TR-dependent processes were disrupted. In addition, we analyzed the regulatory element upstream of human *MSI1* gene using an algorithm-based search for binding sites (NHR scan). We found that the human sequence contained a number of DR4 type response elements, i.e. potential TR binding sites (45–47), as did the mouse.

*Sox2* maintains NSC pluripotency (25), suggesting reduced stem cell renewal, and reduced *Sox10* levels could imply altered fate choice. These results correspond to the well-documented role of THs as a major factor implicated in the regulation of NSC cycling (48), determination of NSC fate towards a neuronal or a glial cell (25, 49) and modulation of vertebrate neurodevelopment (50). Thus, any alteration of TH availability and action on gene expression by 4’-OH-PPF could have direct consequences for neurogenesis and brain morphology (51). Such modifications could also affect craniofacial development as severe hypothyroidism during embryonic development leads to aberrant brain development and craniofacial hypoplasia (26, 27, 52).

### 4’-OH-PPF combined with ZIKV disrupts key neuroprogenitor genes

A series of our results indicated that by itself 4’-OH-PPF already disrupts certain neuroprogenitor genes. In the presence of 4’-OH-PPF, MSI1 levels strongly increased in tadpole brain regions bordering the ventricular system, where the majority of NSCs reside during brain development. This corresponds to what is found in other vertebrates (24). Further, exposing mouse neuroprogenitors to 4’-OH-PPF for 6 days increased levels of *Msi1* and *Klf9*, another T_3_-inducible gene that promotes neuronal development (53, 54) (**Fig S3**). *Msi1* is also a key negatively TH-regulated gene in NSCs (25), and MSI1 is crucial for ZIKV replication in NSCs, interacting with ZIKV RNA (24).

We found that 4’-OH-PPF alters NSC proliferation and the growth of mouse neurospheres, and significantly decreased neuroprogenitor intracellular RNA content with and without ZIKV infection. However, the greatest reduction in intracellular content was seen when the virus and 4’-OH-PPF were both present. Similar results were found in a study using human IPC-based neurospheres, the diameter of which reduced 2-fold following ZIKV infection (8). Our results not only confirmed that 4’-OH-PPF reduced neurosphere numbers, but also had cumulative effects with the virus reflected by the increased numbers of dysregulated genes. We conclude that ZIKV combined with 4’-OH-PPF strongly affects the transcriptional and translation machinery, as seen in the total RNA contents in MSI1-positive cells, reducing survival. Similarly, ZIKV infection induced double-strand breaks in host cells to favor its own replication (55), and increased apoptosis (36). These processes can be directly linked to microcephaly and cortical malformations in ZIKV-infected neonates (56).

Most importantly, most key TH target genes were not dysregulated by 4’-OH-PPF alone, but they were when combined with ZIKV infection. These included *Gfap, Egfr, Nestin, Ng2, Dlx2* and *Dcx*, which are all implicated in NSC self-renewal and neuronal and oligodendroglial commitment, processes underlying normal corticogenesis. These findings offer a plausible mechanism to explain the potential increase of susceptibility to ZIKV in regions where intensive insecticide use was seen.

However, we did not find increased numbers of viral particles per cell, although they remained in the same order of magnitude. However, our neurosphere system has the constraint of being neither a human or a primate-based system, nor an *in vivo* mouse system, for which one could develop a robust 4’-OH-PPF pre-exposure protocol that can be potentially extrapolated to humans. Further, the fetal dose of 4’-OH-PPF nor materno-fetal transmission cannot be perfectly replicated. Nevertheless, work on human neuroprogenitors does show that ZIKV infection attenuates proliferation (7), a result that we also found on mouse neurospheres.

## Conclusions

We have shown that the main metabolite of PPF, 4’-OH-PPF, dysregulates NSC biology by interfering with TH signaling and cell proliferation. These effects are exacerbated when ZIKV co-infection is present. More specifically, the combined effects of 4’-OH-PPF and the virus were seen on key TH target genes required for cortical development. This alone argues for reconsidering the use of the insecticide at its current concentrations in certain Brazilian states, especially given the half-life for PPF in anaerobic conditions can be up to two years (57). Even though Europe is not proposing to use PPF at these concentrations, it has recently been reauthorized (58). The data provide another example of how a drug or a pesticide can exacerbate the outcome of viral infection.

## 4. Materials and methods

### 4.1 *In vitro* luciferase assay

JEG3-cell culture transfections were performed as previously described (59). Briefly, FLAG-tagged human TRα1 was overexpressed in JEG3 cells together with a DR4-TRE luciferase reporter construct and pMaxGFP as a transfection control. After 24 h, cells were incubated for 24 h in DMEM/F12 + 0.1% BSA containing 1 nM of T_3_ and the indicated concentrations of 4’-OH-PPF or the TRα1-antagonist, NH3. Luciferase activity was measured in cell lysates as previously described (60). The ratio of the luciferase and GFP activities was calculated to correct for transfection efficiency. The results are expressed as percentage of the ratio at 0 nM 4’-OH-PPF or NH3. Two independent experiments were performed in triplicate.

### 4.2 In vivo studies in *Xenopus laevis*

#### XETA assay

We used modified Xenopus Euletheroembryo Thyroid Assay (31) to assess the thyroid-disrupting potential of environmental concentrations of 4’-OH-PPF *in vivo*. This assay used the Tg(*thibz*:eGFP) transgenic *Xenopus laevis* tadpoles.

#### Chemical exposure

The stock solutions 4’-OH-PPF (HPC – Ref. 675366) were prepared according to following protocol: 15 mg 4’-OH-PPF were dissolved in 5 ml of Dimethyl sulfoxide (DMSO, Sigma - CAS 67-68-5) to create a 3 g/L stock solution (stored at -20 °C). The final exposure solution was prepared daily prior to exposure using fresh aliquots of stock solutions (prepared by cascade dilutions from 3 g/L stock) by adding 1 μL of stock solution to 10 mL of Evian water. All groups including control containing 0.01% DMSO. Due to the rapid conversion rate of PPF to 4’-OH-PPF in the organism (21), we used maximum concentrations of 4’-OH-PPF in drinking water according to WHO recommendations. Fifteen NF 45 Tg(*thibz*:eGFP) *Xenopus laevis* tadpoles were placed into each well of a 6-well plate. Each well corresponded to one exposure group. Eight mL of previously prepared final exposure solution was added per well. Tadpoles were exposed for 72 h at 23 °C in dark with daily renewal at the same hour for the XETA assay. For brain gene expression analysis, exposure was limited to 24 h. Tadpoles were exposed to 4’-OH-PPF ± 5 nM T_3_ (3,3′,5-triiodo-L-thyronine sodium salt (T_3_) (Sigma - CAS 55-06-1)).

#### GFP signal imaging

At the end of exposure, tadpoles were rinsed in Evian and anaesthetized using Ethyl 3-aminobenzoate methanesulfonate salt (MS 222) (Sigma - CAS 886-86-2, 100 mg/L; Stock 1 g/L - 1 g of MS-222 and 1 g NaHCO_3_ dissolved in 1 L of Evian, pH adjusted to 7.4-8; stock diluted 10x in Evian). Once anaesthetized, one Tg(TH/*bzip*:eGFP) tadpole was placed per well into 96-well plate (black, conic based) and positioned using a fine transfer pipette so the ventral region of the tadpole was facing upwards. To acquire color images, we used an Olympus SZX12 Stereo Microscope equipped with a 1.2X objective and a 25X magnification used, long pass GFP filters and a QImaging Exi Aqua CCD camera. Images were captured using the program QCapture Pro 6.0 (QImaging) with 3 s exposure time. After the image acquisition, tadpoles were euthanized in MS-222 1 g/L, fixed overnight (ON) in paraformaldehyde 4% and stored at -20 °C in cryoprotectant (150 g of sucrose (Sigma - CAS 57-50-1) and 150 mL of ethylene glycol (Sigma - CAS 107-21-1) volume adjusted to 500 mL by PBS 1x). Using the program ImageJ, non-specific signal was excluded by splitting the image into three layers (red, blue and green channel) followed by subtraction of the red and blue channel from the green one. Integrated density of the green channel was then quantified.

#### RT-qPCR

At the end of the 24 h exposure window, tadpoles were rinsed and anaesthetized in 100 mg/L MS-222 (prepared as in 2.1.3). RNA-free dissecting tools were used to dissect tadpole brains. Two brains were placed in one microdissection tube containing 100 μL lysis solution from an RNA extraction kit (Ambion RNAqueous). Tubes were snap-frozen and stored at -80 °C prior to RNA extraction according to the manufacturer’s instructions (Invitrogen). The quality of each sample was verified using Agilent bio-analyzer. cDNA was synthetized using Reverse Transcription Master Mix (Fluidigm). The synthetized cDNA was 20x diluted (5 μL of DNA in 95 μL of nuclease free water). qPCR mix contained following components: 0.15 μL of reverse primer (10 pM), 0.15 μL of forward primer (10 pM) (**Table 1**), 1.7 μL of nuclease free water and 3 μL of Power SYBR master mix per reaction. For the final reaction, 5 μL of previously prepared qPCR mix was added in each well of 384 well plate together with 1 μL of diluted cDNA. Every reaction was performed in duplicate. Water controls and RT-controls were included on every plate. Comparative Ct measurements including the melting curve were quantified using QuantStudio 6 flex qPCR device (Life technologies). The geometric mean of two housekeeping genes (*ef1a* and *odc*) of each sample was quantified and excluded from the Ct values corresponding to the sample (resulting in ΔCt value). Median values of the control group of each gene were calculated and extracted from each ΔCt value to normalize, providing ΔΔCt values. Fold change was calculated using formula =(power 2; - ΔΔCt value of a sample).

**Table 1:**
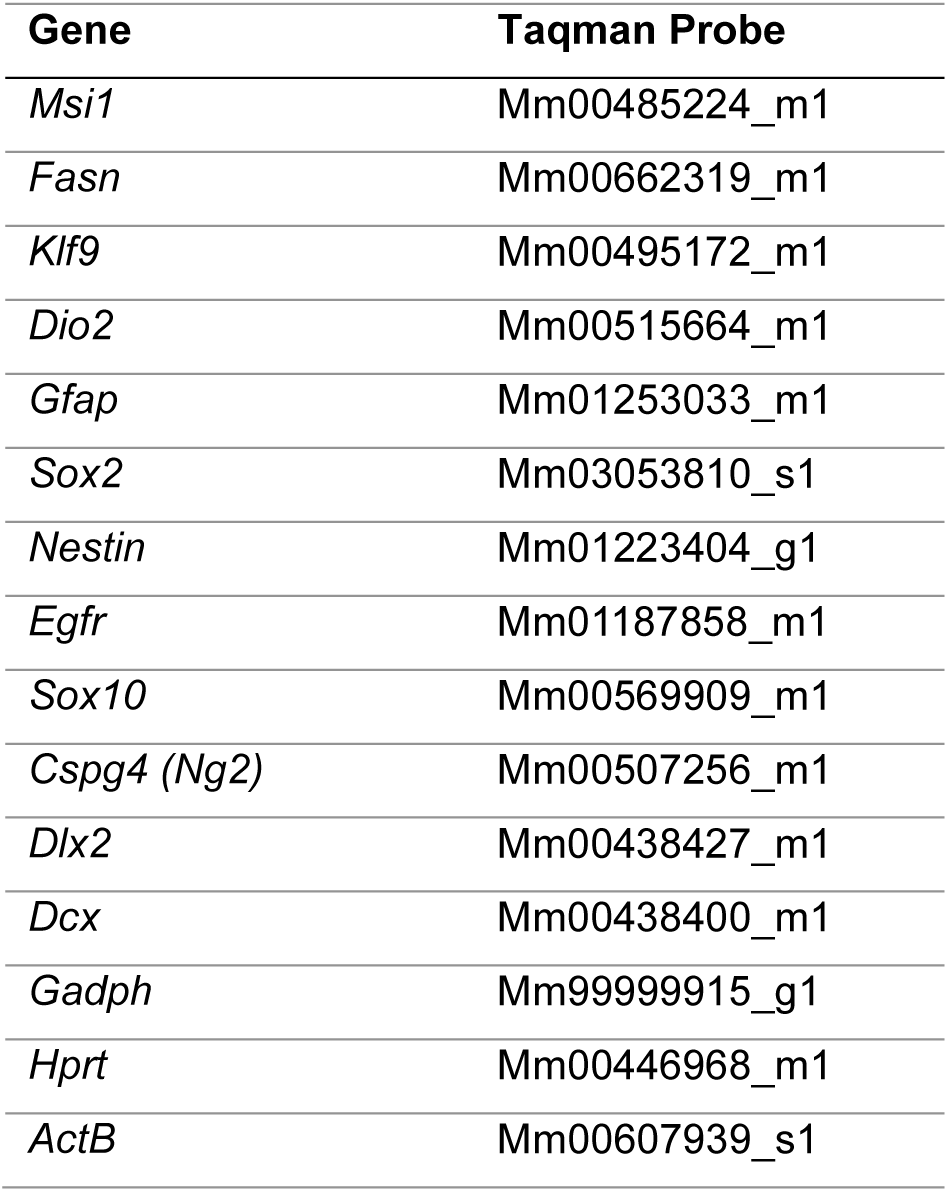
Specific Taqman probes used for qPCR experiments.

| **Gene** | **Taqman Probe** |
| --- | --- |
| *Msi1* | Mm00485224_m1 |
| *Fasn* | Mm00662319_m1 |
| *Klf9* | Mm00495172_m1 |
| *Dio2* | Mm00515664_m1 |
| *Gfap* | Mm01253033_m1 |
| *Sox2* | Mm03053810_s1 |
| *Nestin* | Mm01223404_g1 |
| *Egfr* | Mm01187858_m1 |
| *Sox10* | Mm00569909_m1 |
| *Cspg4 (Ng2)* | Mm00507256_m1 |
| *Dlx2* | Mm00438427_m1 |
| *Dcx* | Mm00438400_m1 |
| *Gadph* | Mm99999915_g1 |
| *Hprt* | Mm00446968_m1 |
| *ActB* | Mm00607939_s1 |

#### Immunohistochemistry

After 24 h of exposure, tadpoles were euthanized using MS-222 (1 g/L) and fixed in 4% paraformaldehyde (Sigma - CAS 30525-89-4) for 3 h at room temperature (RT). After fixation, the tadpole brains were dissected and placed in PBT (1% Triton X-100 in PBS) for at least 18 h at 4 °C. Next, the brains were blocked in 10% (vol/vol) normal goat serum (NGS) in PBT for 3 h at 4 °C. The primary antibody (Musashi-1, polyclonal rabbit: Abcam ab21628, UK) was diluted 1/300 in 10% NGS in PBT and the brains were incubated in the solution ON at 4 °C. Next morning, the plates were removed to RT for 1 h prior to multiple washes in PBT for at least 8 h. The secondary anti-rabbit antibody Alexa Fluor 488-conjugated (Thermofisher Scientific) diluted 1/400 in 10% NGS in PBT was applied ON at 4 °C. Brains were then washed several times in PBS for 4 h (RT). Images of the dorsal side of the brains were taken using a Leica MZ16F stereomicroscope equipped by QImaging Retiga-SRV camera. The mean integrated density of each image was calculated with the threshold set to 60 using ImageJ software.

#### Tadpole behavioral assays

After 72 h exposure, tadpoles were rinsed in Evian water and placed in 12-well plates with one tadpole per well each containing 4 mL Evian water. Each group was given 10 minutes acclimatization time, prior to the trial. Tadpole movements were tracked for 10 minutes with 30 s light/30 s dark intervals using a Daniovision (Noldus) device. Total distance travelled during each 10 s of the 10 minute trials were quantified and exported using the Ethiovision XT 1.5 program. All values were normalized to the mean of the control group of the [0 s-10 s] time period.

#### Structural measurements of head and brain compartments

Tadpoles previously fixed in 4% paraformaldehyde during 3 h at RT were placed in a Petri dish and manually positioned to expose the dorsal part of the body. Color images of tadpole heads on black background were taken using a Leica MZ16F stereomicroscope equipped by QImaging Retiga-SRV camera. The area of the brain and head and the width of forebrain, midbrain and forebrain/midbrain junction were measured in ImageJ with manually defined ROIs and the head/brain ratio was calculated.

### 4.3 In vitro mouse neurosphere assays

#### Animals

C57BL/6JRj 8 week-old wild-type male mice were purchased from Janvier Labs (Le Genest St. Isle, France) and kept under standard conditions (22 °C, 50% humidity, 12 h light/dark cycle) with food and water *ad libitum*. Animals were handled and cared for in accordance with the EU Directive of 22 September 2010 (2010/63/EU). All procedures were conducted according to the principles and procedures in Guidelines for Care and Use of Laboratory Animals and validated by local and national ethical committees (CEEA Cuvier n°68).

#### Virus

Virus strain H/PF/13 from EVA program (ref 23484; sequence I.D. KJ776791.1) was amplified *in vitro* in a single round on C6/36 mosquito cells (ATCC® CRL-1660™) and titrated using VERO E6 cell line (ATCC CRL-1586) and by QRT-PCR with a titer of 2 10^7^ TCID50%/mL corresponding to around 5.4 10^8^ RNA genome copies/mL. The virus stock is aliquoted and stored à -80 °C until use.

#### Adult mouse neurosphere assay

Five mice were sacrificed per neurosphere culture. Brains were removed and the lateral SVZs were dissected in cold DMEM-F12-glutamax 1/50 Glc 45% and subsequently incubated in digestion medium (papain [Worthington], DNase [Worthington], L-cysteine [Sigma-Aldrich]) for 30 min at 37 °C. Cells were dissociated every 10 min to obtain a single cell suspension. Cells were then spinned down (1000 rcf, 5 min) and equally distributed in wells containing growth medium (DMEM-F12-glutamax [Gibco], 40 μg/mL insulin [Sigma], 1/200 B-27 supplement [Gibco], 1/100 N-2 supplement [Gibco], 0.3% glucose, 5 mM Hepes, 100 U/ml penicillin/streptomycin) containing the growth factors (Gfs) EGF and FGF2 (20 ng/mL [Peprotech]). Using a radio-immunoassay, we found that the B-27 medium contains 0.3 nM T_3_.

In a first experiment, 10^4^ cells/well in 100 µL/well were exposed in 96-well plates (12 wells/group). On day 0, 10 nM T_3_ (Sigma - CAS 55-06-1) together with 10^−2^, 10^−1^, or 3×10^−1^ mg/L 4’-OH-PPF was applied for seven days in a 5% CO_2_, 20% O_2_ environment at 37 °C. Thirty µL/well of the growth medium was supplemented every two days. At day 7, brightfield images of neurospheres were taken (ZEISS AXIO Zoom V16 macroscope; 21 images/well) and the projected areas of neurospheres were measured using the ImageJ software. Then, after 5 min centrifugation at 1000 rcf, cell pellets were frozen at –20 °C until RNA extraction.

In a second experiment, 3×10^5^ cells in 5 ml of medium were exposed with 0.01% DMSO ± 4’-OH-PPF 10^−1^ mg/L from day 2 to day 7 of *in vitro* proliferation, with 1.5 ml of this medium supplemented every other day. At day 7, the neurospheres were dissociated and infected with a fresh aliquot of ZIKV stock thawed at 37°C and diluted in DMEM medium to a multiplicity of infection of 0.5 particles per cell during 1 h at 37 °C (i.e. ± 1 million cells in 80 µl of viral suspension). The cells were then extensively washed (5 times with 3 mL of DMEM) before they were distributed in 6 wells per conditions (see figure 3A). Six replicates of 150 000 cells/well for the control (0.01% DMSO) and 4’-OH-PPF condition were used. Gfs (EGF, FGF2: 20 ng/mL) ± 10^−1^ mg/L 4’-OH-PPF were added every other day. Viral replication in differentiated neurospheres was analyzed every day for a period of 4 days by RT-PCR using primers and probes derived from (61) (ZIKV_F, and ZIKV_R) encompassing a small segment coding for the E protein. Briefly, 5 µL of culture supernatants from each well and time point were diluted 1/100. Then, 3 µL of the dilutions were mixed with the QRT-PCR medium containing primers, probes, enzyme and buffer (Supersript III platinum one step qPCR system from Invitrogen/ThermoFisher Scientific) within a 96 well plate and PCR ran on a Bio-Rad CFX thermocycler. Results were quantified relative to ZIKV Vero supernatant diluted from 10^7^ RNAcopies/mL to 330 copies/mL that was previously calibrated as described in (62). At day 4, cells were spinned down 5 min at 1000 rcf and frozen at -20 °C until RNA extraction.

#### Gene expression analysis

After 7 days of proliferation, neurospheres were pooled per condition, RNA was extracted (ThermoFisher RNAqueous MicroKit) and retro-transcriptions were performed (Fluidigm) following manufacturers’ instructions. Pre-amplifications (10 cycles) and qPCR were performed using TaqMan Preamp MasterMix and TaqMan Universal PCR MasterMix, respectively (ThermoFisher), and specific TaqMan probes. Every qPCR amplification was run in triplicate. ΔCt values were calculated between target genes and the geometrical mean of three housekeeping genes, *ActB, Gapdh* and *Hprt*. The fold change in target gene expression was performed as described above.

#### Immunostaining of Musashi-1

Neurospheres were dissociated into single cells and 50.10^3^ cells were plated onto 24-well Glass Slips (Millicell EZSLIDE, PEZGS0896) coated with 0.1mg/ml Poly-D-Lysine (Sigma-Aldrich), in culture medium without EGF/FGF2 for 1 h. Cells were fixed with 4% paraformaldehyde for 10 min at RT and blocked with 10% donkey serum, 0.2% triton and 1% BSA in PBS for 30 min at RT. Cells were then incubated with primary antibody (anti-Musashi-1: 1/300, polyclonal rabbit: Abcam ab21628, UK) in blocking solution for 48 h at 4 °C. After three 5 min washes in PBS at RT, cells were incubated with a fluorescent secondary antibody (donkey anti-rabbit Alexa Fluor 488-conjugated, Invitrogen, 1/500) in 1% donkey serum, 0.2% triton and 1% BSA in PBS for 2h at RT. Cells were washed three times in PBS, incubated with DAPI for 5 min at RT and mounted with Prolong Gold antifade reagent. Fluorescence images (10 images per condition) were acquired using a Leica TCS-SP5 confocal microscope and processed using the ImageJ software. Quantifications have been done on Max Intensity Z projection of 4 µm stack images.

#### 4.3. Statistical analysis

Statistical analysis was performed using GraphPad Prism v7.00. The D’Agostino & Pearson normality test was done to determine if values follow Gaussian distribution (parametric) or not (non-parametric). One-way ANOVA was done in the case of parametric distribution and followed by a Tukey’s or Dunnett’s multiple comparison post-test. Two-way ANOVA followed by Bonferroni’s multiple comparisons test was used to analyse ZIKV replication. A Kruskal-Wallis test with Mann-Whitney multiple comparison post-tests was used for non-parametric data. For comparison of two groups, a t-test (parametric) or a Mann-Whitney U test (non-parametric) was used. Data are presented as scatter plots depicting mean ± standard error of mean (SEM) or standard deviation (SD). P values <0.05 were thereby considered as statistically significant.

## 5. Acknowledgments and funding sources

We are grateful for the work in the Xenopus facility of Gérard Benisti, Philippe Durand and Jean-Paul Chaumeil and the ZIKV QRT-PCR were performed in the L2I platform (head N Bosquet) of IDMIT/ CEA/INSERM UMR1184 by Marco Leonec. We also thank the ImagoSeine plateform of the Institut Jacques Monod (Université René Diderot) and the CeMIM plateform of the Muséum National d’Histoire Naturelle (MNHN). We would like to thank all our colleagues for their comments on the manuscript, and Fiene Kuijper for analysis of the mouse neurosphere data. We are grateful for the work of Theo Visser. Pieter Vancamp was supported by a basic science grant from the European Thyroid Association, and by the Fondation pour la Recherche Médicale (FRM, SPF201909009111). This work was supported by grants from Centre National de la Recherche Scientifique (CNRS), Muséum National d’Histoire Naturelle (MNHN), PNREST THYPEST EST-2014-122 and European Union contract EDC MIX RISK_GA N°634880. PR Zika work was supported by the EU H2020 program ZIKAlliance GA N°734548.

---

**Supplementary figure 1.**
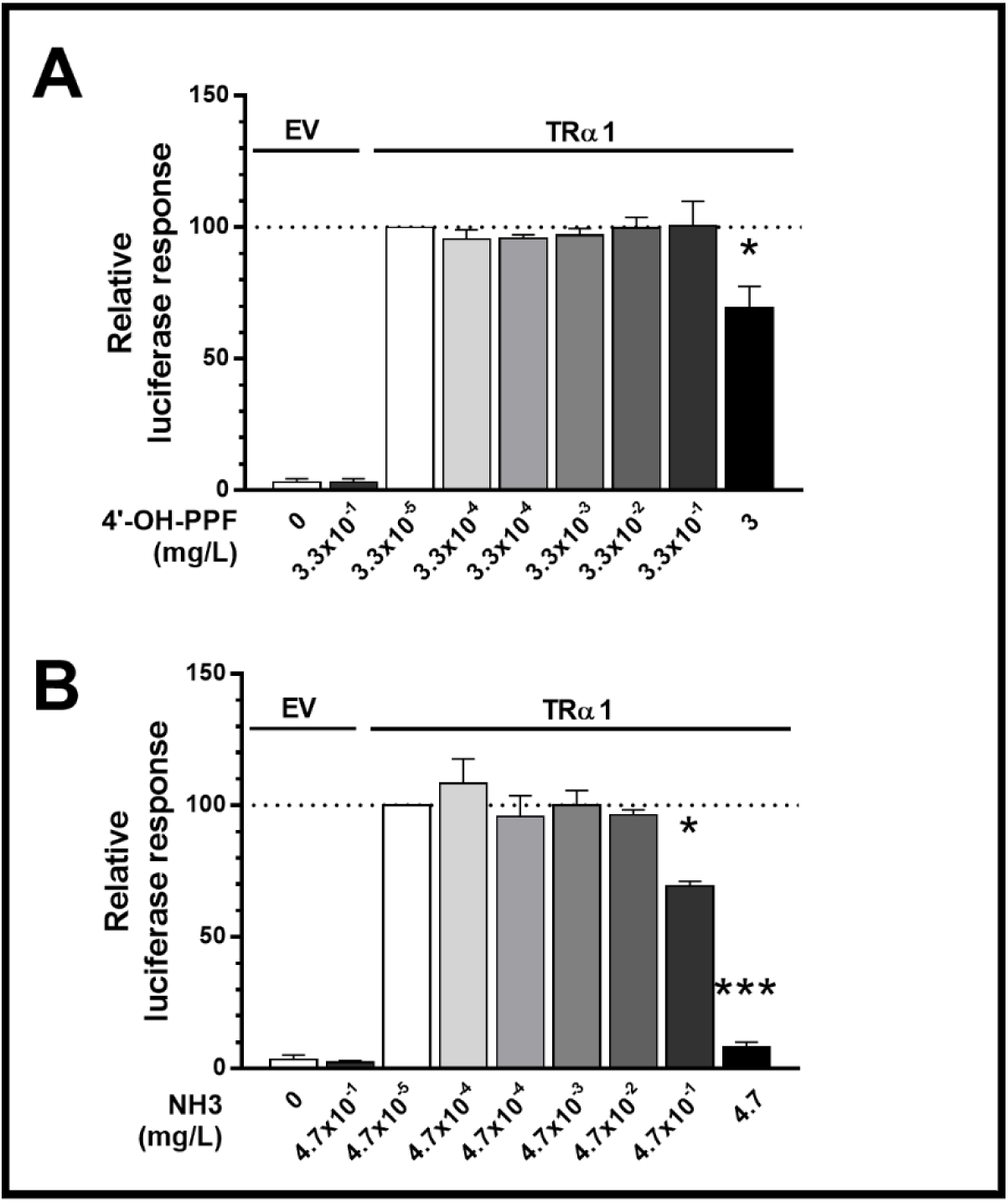
4’-OH-PPF acts as TRα1 antagonist. (**A, B**) The transcriptional activity of TRα1 stimulated by 1 nM T_3_ was significantly reduced under increasing concentrations of 4’-OH-PPF (**A**) and NH3 (**B**), the latter being a strong TH antagonist. The data are presented as mean ± SEM of two independent experiments performed in triplicate (One-way ANOVA p<0.05; Tukey’s post-test compared to 0 nM competitor, *p<0.05, ***p<0.001). EV: empty vector control.

**Supplementary figure 2.**
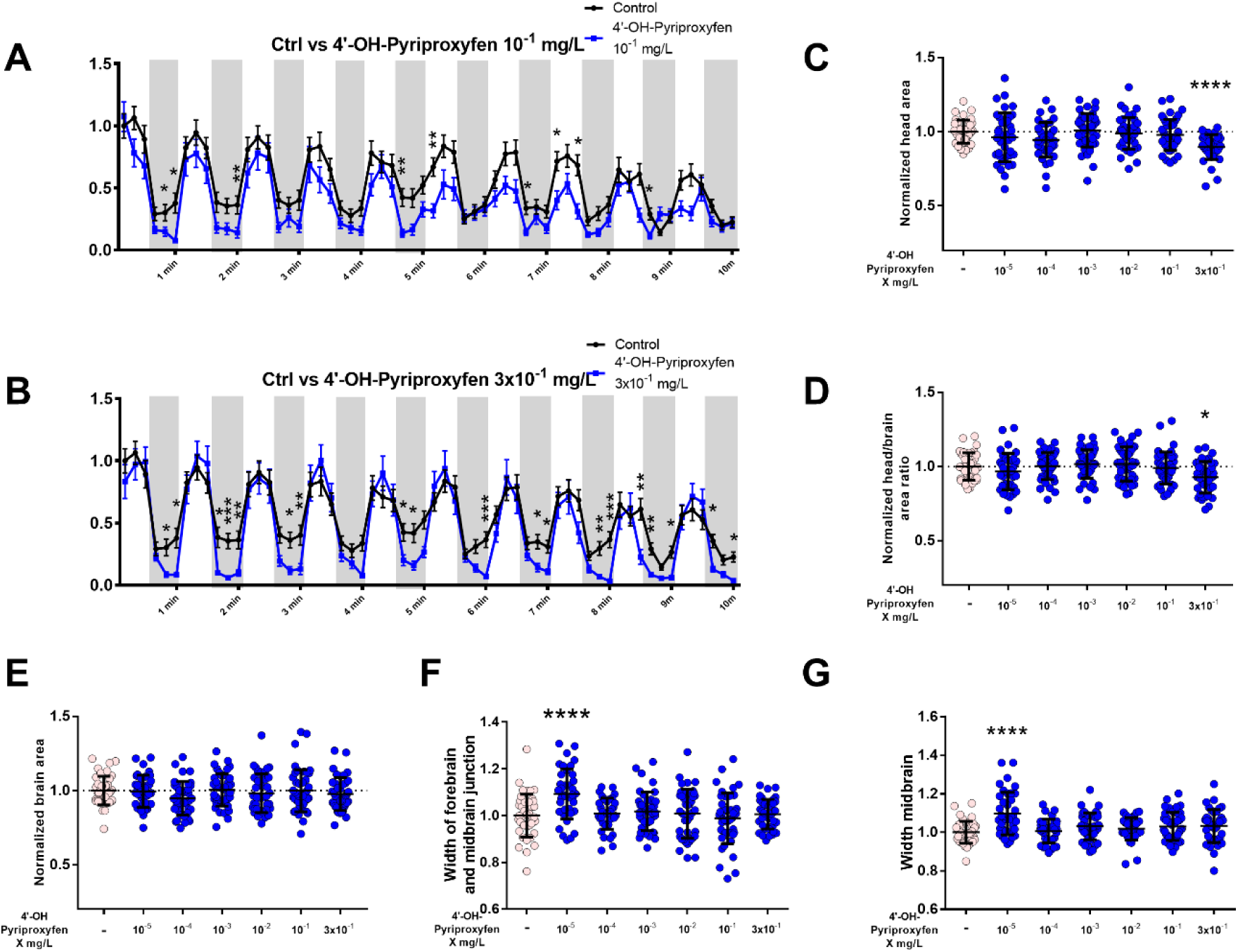
Exposure to 4’-OH-PPF reduces tadpole mobility and the lowest dose increases fore- and midbrain width. **(A-B)** Total distance travelled by tadpoles exposed to 10^−1^ g/mL 4’-OH-PPF (**A**) or 3×10^−1^ g/mL 4’-OH-PPF (**B**) during a 10 minute trial, with distance travelled recorded every 10 s. Dark backgrounds represent unlit periods, white backgrounds represent light periods. Values were normalized to the first 10 s period of the control group (Pool of 3 independent experiments; n = 12 per experiment, Kruskal-Wallis, mean ± SEM, * p<0.05, ** p<0.01, *** p<0.001). Graphs showing **(C)** normalized head area, **(D)** head/brain area ratio, **(E)** brain area, **(F)** width of fore- and midbrain junction and **(G)** width of midbrain of NF45 tadpoles exposed to 10^−1^ g/mL 4’-OH-PPF for 72 h. Values normalized to the control group (Pool of 3 independent experiments; n=15 per experiment, One-way ANOVA with Dunn’s post-test, mean ± SDs, * p<0.05, **** p<0.0001).

**Supplementary figure 3.**
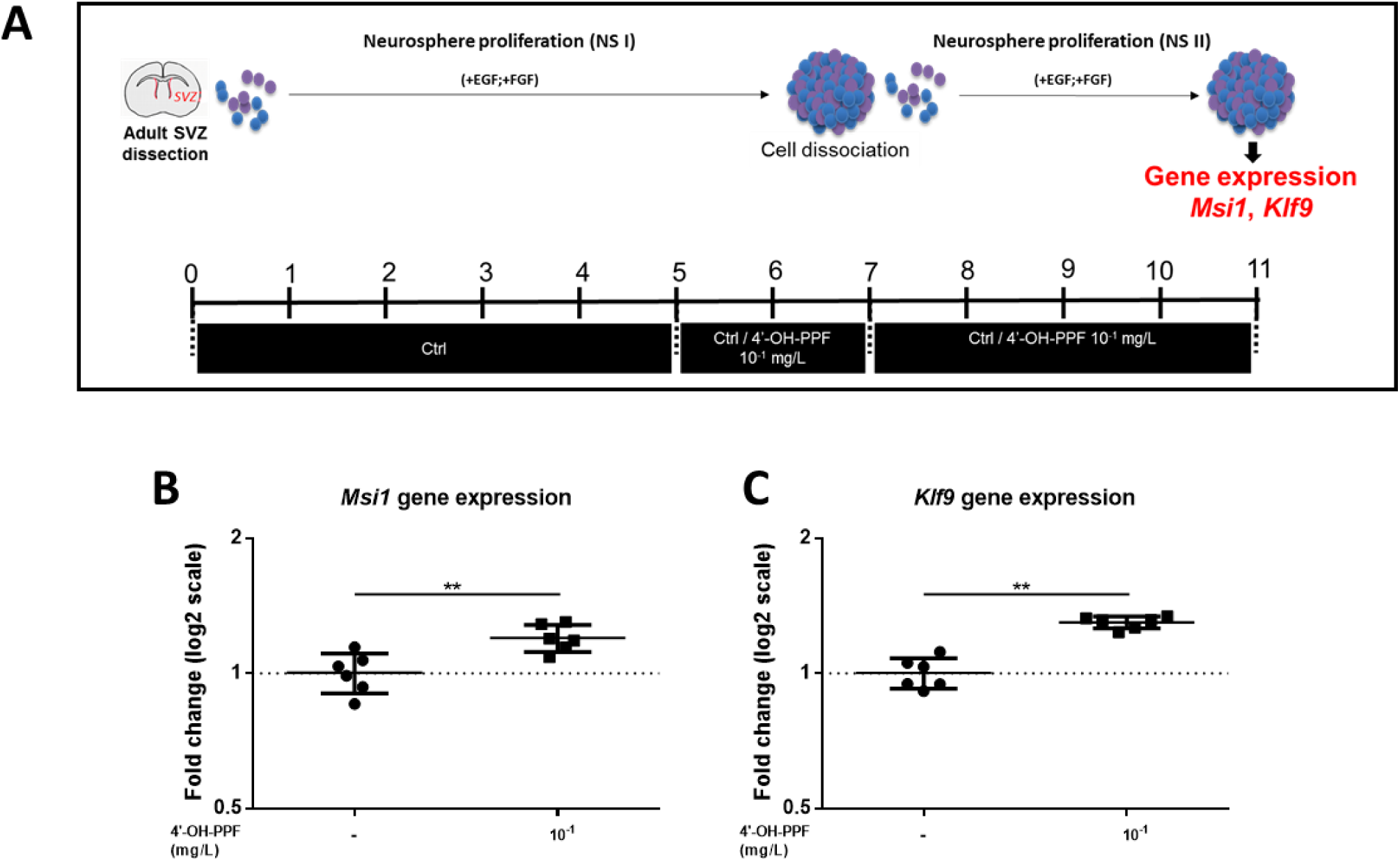
A 6-day exposure to 10^−1^ mg/L 4’-OH-PPF induces *Msi1* and *Klf9* expression in mouse neurospheres. **(A)** Schematic timeline illustrating the experimental design. **(B-C)** Gene expression profile for TH targets *Msi1* **(B)** and *Klf9* **(C)**. Graphs showing mean ± SDs (n=6, Mann-Whitney test, *Msi1* **: p=0.0087, *Klf9* **: p=0.0022).

