## Supplementary figures + legends for "ZIKA virus effects on neuroprogenitors are exacerbated by the main pyriproxyfen metabolite via thyroid hormone signaling disruption"

**Supplementary figure 1**

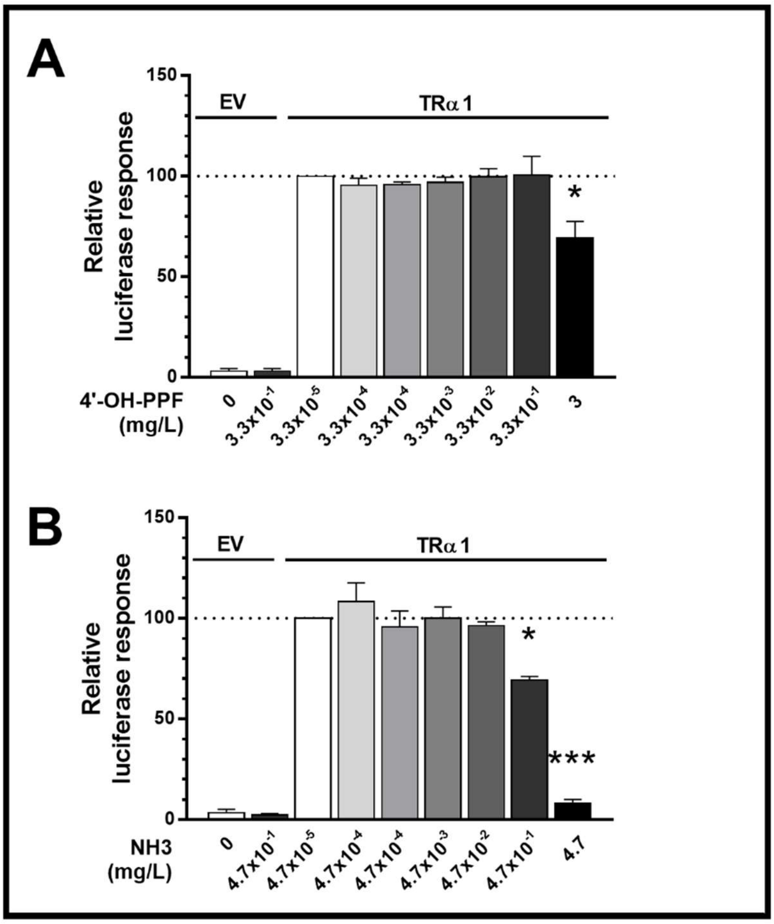


**4’-OH-PPF acts as TRα1 antagonist with an affinity 50x lower than NH3.** (**A, B**) The transcriptional activity of TRα1 stimulated by 1 nM $T_3$ was significantly reduced under increasing concentrations of 4’-OH-PPF (**A**) and NH3 (**B**), the latter being a strong TH antagonist. The data are presented as mean ± SEM of two independent experiments performed in triplicate (One-way ANOVA $p<0.05$; Tukey’s post-test compared to 0 nM competitor, *$p<0.05$, ***$p<0.001$). EV: empty vector control

**Supplementary figure 2**

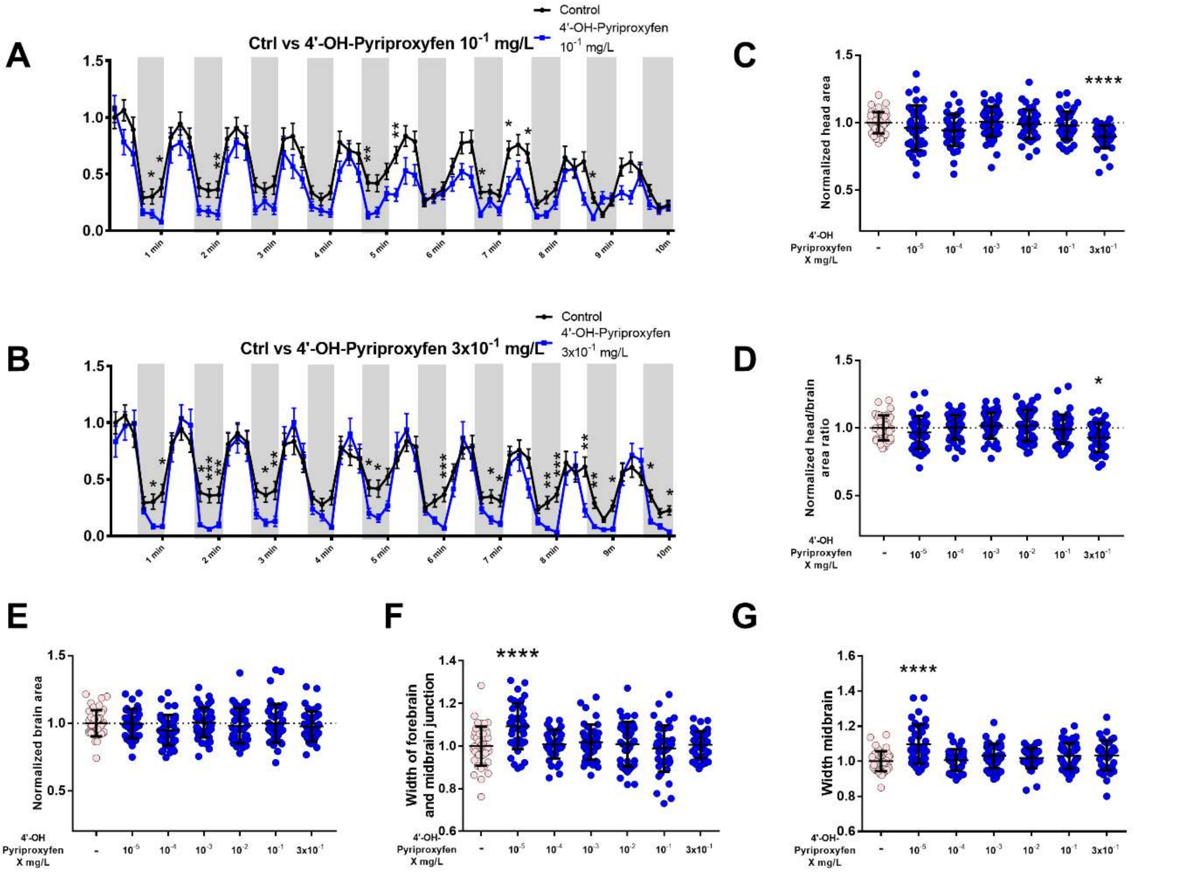


**Exposure to 4'-OH-PPF reduces tadpole mobility and the lowest dose increases fore- and midbrain width. (A-B)** Total distance travelled by tadpoles exposed to $10^{-1}$ g/mL 4'-OH-PPF (**A**) or $3\times10^{-1}$ g/mL 4'-OH-PPF (**B**) during a 10 minute trial, with distance counted every 10 s. Dark backgrounds represents dark periods, white backgrounds represents light periods. Values were normalized to the first 10 s period of the control group (Pool of 3 independent experiments; n = 12 per experiment, Kruskal-Wallis, Mean ± SEM, * $p<0.05$, ** $p<0.01$, *** $p<0.001$). Graphs showing **(C)** normalized head area, **(D)** head/brain area ratio, **(E)** brain area, **(F)** width of fore- and midbrain junction and **(G)** width of midbrain of NF45 tadpoles exposed to $10^{-1}$ g/mL 4'-OH-PPF for 72 h. Values normalized to the control group (Pool of 3 independent experiments; n=15 per experiment, One-way ANOVA with Dunn's post-test, Mean ± SDs, * $p<0.05$, **** $p<0.0001$).

**Supplementary figure 3**

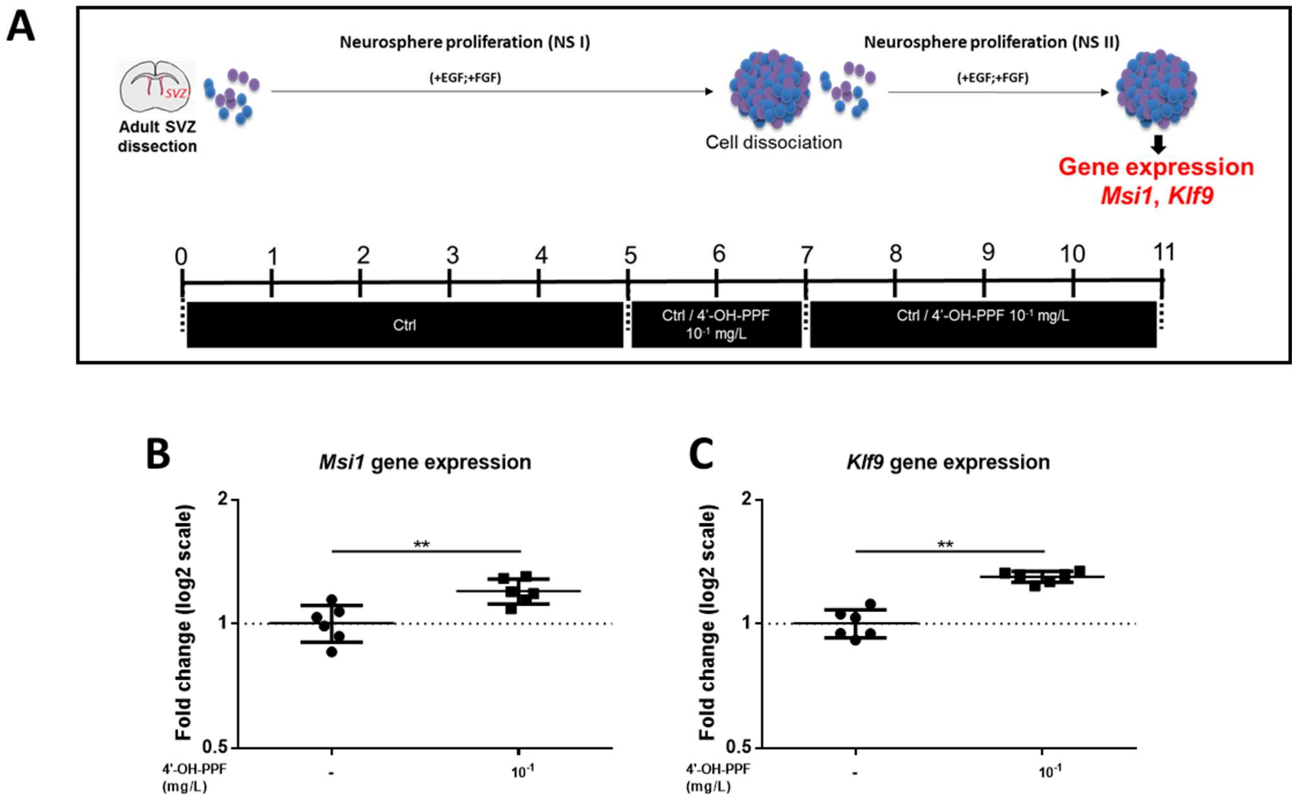


**A 6-day exposure to $10^{-1}$ mg/L 4'-OH-PPF induces *Msi1* and *Klf9* expression in mouse neurospheres. (A)** Schematic timeline illustrating the experimental design. **(B-C)** Gene expression profile for TH targets *Msi1* **(B)** and *Klf9* **(C)**. Graphs showing mean ± SDs (n=6, Mann-Whitney test, *Msi1* **: p=0.0087, *Klf9* **: p=0.0022).
